# Modelling Inflammatory Endothelial Dysfunction: A Human In Vitro Platform for Translational Research

**DOI:** 10.1101/2023.10.31.564901

**Authors:** Maria Cheremkhina, Aaron Babendreyer, Christopher T. Neullens, Susanne Krapp, Alessa Pabst, Kim Ohl, Stephan Ruetten, Andreas Ludwig, Christian G. Cornelissen, Stefan Jockenhoevel, Klaus Tenbrock, Anja Lena Thiebes

## Abstract

Systemic inflammation presents a significant challenge to the long-term function of biohybrid implants. While endothelialisation of biohybrid implants has been shown to improve device hemocompatibility, its feasibility under the influence of patients’ inflammatory status remains largely unexplored. To investigate this, we developed a controlled *in vitro* model which allows to study endothelial dysfunction under inflammatory stress. Endothelial cells were cultured on polydimethylsiloxane under physiological shear stress and exposed to lipopolysaccharide (LPS)-activated peripheral blood mononuclear cells (PBMCs), simulating inflammatory conditions. Endothelial morphology and confluence was assessed using immunohistochemistry and scanning electron microscopy. Leukocyte adhesion was evaluated directly as well as indirectly, using flow cytometry to analyse cell adhesion molecules. Quantitative PCR was used for gene expression analysis of inflammatory mediators. Notably, neither LPS nor PBMCs alone induced endothelial disruption, whereas their combination significantly impaired endothelial confluence: Inflammatory activation led to substantial loss of endothelial confluence, increased leukocyte adhesion, and elevated expression of adhesion molecules ICAM-1, VCAM-1, and E-selectin. Gene expression analysis highlights the upregulation of inflammatory mediators, such as IL-6, IL-8, IL-10, and MCP-1. This study underscores the challenges of implementing endothelialisation in biohybrid devices, particularly in patients with systemic inflammation. By considering translational hurdles, this work contributes to the development of clinically viable biohybrid constructs and highlights the importance of considering inflammatory dynamics when designing next-generation implants.

## Background

Biohybrid implants are a promising strategy to improve hemocompatibility of blood-contacting devices. One of the strategies to create a biohybrid device is the endothelialisation of its artificial surfaces [1, 2]. This approach has been applied to several devices, such as vascular grafts and gas exchange membranes of oxygenators [3, 4]. General feasibility of endothelialisation has been demonstrated under standard cell culture conditions *in vitro* for example, for oxygenator membrane [5, 6]. Nevertheless, these conditions do not adequately replicate the complex inflammatory state of many patients. High levels of inflammatory cytokines in blood are characteristic of patients undergoing extracorporeal membrane oxygenation (ECMO) treatment, as respiratory failure is often associated with systemic inflammation, whether in acute lung failure or chronic obstructive pulmonary disease (COPD) [7, 8]. Moreover, it has been reported that ECMO treatment itself can induce systemic inflammatory response syndrome (SIRS), which leads to increased mortality [9, 10].

The behaviour of endothelial cells under inflammatory conditions *in vivo* has been extensively investigated [11, 12]. While endothelial barrier function is maintained in healthy conditions by endothelial cell glycocalyx and intercellular junctions, increased cytokine production during inflammation causes upregulation of adhesion molecules, which in turn allows for leukocyte adhesion and transmigration [13, 14]. This process, while crucial for tissue healing, disrupts intercellular junctions, allowing leukocyte migration to inflamed tissue [15]. In the context of endothelialized biohybrid implants, this inflammatory response poses a significant challenge: the disruption of intercellular junctions can expose the artificial surface to blood, increasing device thrombogenicity and undermining the biohybrid implant concept.

In the current study, we evaluate the behaviour of endothelial cells on a polydimethylsiloxane (PDMS) gas exchange membrane *in vitro* under inflammatory conditions. Since flow conditions can significantly alter endothelial cell behaviour [16], cells are conditioned with relevant wall shear stresses in a microfluidic system. We investigated endothelial cell layer integrity, expression of adhesion molecules and leukocyte adhesion as well as changes in gene expression in standard culture and inflammatory conditions. Our goal is to better predict endothelial cell behaviour in biohybrid devices during future clinical application.

## Materials and Methods

### Isolation and Culture of Human Umbilical Vein Endothelial Cells

Human umbilical vein endothelial cells (HUVECs) were isolated from umbilical cords according to an established protocol [17]. Umbilical cords were provided by the Centralized Biomaterial Bank of the RWTH Aachen University (cBMB) according to its regulations, following RWTH Aachen University, Medical Faculty Ethics Committee (cBMB project number 323) and the Department of Gynaecology and Perinatal Medicine (Univ.-Prof. Dr. Stickeler). The patients’ authorized representative provided informed consent. Culture of HUVECs was performed in endothelial cell growth medium 2 (EGM2, Promocell) with 1% antibiotic-antimycotic solution (ABM, Thermo Fischer) in gelatine-coated (2%, Sigma-Aldrich) tissue-culture flasks (Greiner) in a humidified incubator at 37 °C and 5% CO_2_. Characterization of HUVECs was performed by flow cytometry analysis for CD31 (PECAM-1), CD105, CD144, CD146, and CD90. As shown in the Supplementary Table S1, HUVECs were positive for CD31, CD105, CD144, and CD146, and negative for CD90, which confirmed their endothelial cell characteristics. HUVECs (*n = 3* independent donors) were used in passage 4 for all experiments.

### Isolation of Peripheral Blood Mononuclear Cells

Peripheral blood mononuclear cells (PBMCs) were isolated from freshly drawn venous blood. Blood was obtained from healthy male donors between 25 and 35 years of age after approval by the local ethics committee of the Medical Faculty of RWTH Aachen University Hospital (EK 22-390) and providing informed consent. Briefly, a Pancoll gradient (Pan-Biotech) was used to separate PBMCs from other blood components. Erythrocytes were lysed according to manufacturer‘s instructions in erythrocyte lysis buffer (Pan-Biotech). PBMCs were used directly after isolation for all experiments.

### Static Culture of Endothelial Cells on Gas Exchange Membranes for Thrombin Generation Assay

Polydimethylsiloxane (PDMS) membranes (800 µm, Limitless Shielding) were washed with soap, absolute ethanol (70%, Merck), and ultra-pure water (Sartorius). Circular samples (6 mm diameter) were glued (Elastosil^®^ E50 N Transparent silicon rubber, Wacker Chemie AG) to the bottom of 24-well plates (Greiner). PDMS membranes were functionalised with RGD peptides (a binding motif for cellular integrins) as previously described [18].

HUVECs were resuspended in medium and seeded with a concentration of 7 × 10^5^ cells/cm^2^. Endothelial cells were incubated for 24 h to allow cell attachment. The membranes were checked for full confluence before performing the thrombin generation assay.

To evaluate the thrombogenicity of the biofunctionalized and endothelialised membranes, a thrombin generation assay (TGA, Haemoscan) was performed. The membranes were carefully washed with DPBS, and the assay was performed according to the manufacturer’s instructions compliant with ISO 10993-4:2017. Briefly, all samples were incubated with human plasma before adding the TGA reagents. Thrombin generation was measured at 1, 2, 3, and 4 min after reagent addition. The optical density was measured at 405 nm and 540 nm with a microplate reader (Infinite^®^ 200Pro, Tecan). Thrombin concentration was calculated from acquired calibration curves. The calculation of maximum thrombin generation was performed according to the manufacturer‘s instructions. Briefly, the thrombin generation between 3 and 4 min was normalized to the membrane surface area, as the highest increase in generated thrombin occurred during this time.

### Dynamic Culture of Endothelial Cells on Gas Exchange Membranes

PDMS membranes were washed as described above, followed by gluing of the membranes to the microfluidic devices (sticky Slides 0.2 Luer, Ibidi) with 250 µm channel height as previously described [19] and coated with RGD. For endothelialisation, HUVECs were resuspended in EGM2 medium with 1% ABM at a concentration of 5 × 10^6^ cells/mL and seeded in the microfluidic channels. Microfluidic devices with seeded cells were incubated statically for at least 2.5 h until full cell attachment while medium exchange was carefully performed every 30 minutes. Afterwards, three microfluidic devices were connected in series to the bioreactor system as depicted in Supplementary Figure S1. Laminar unidirectional flow of 8.4 mL/min was applied to the cells in microfluidic devices, resulting in a wall shear stress (WSS) of 20 dyn/cm^2^. After 24 h of endothelial cell dynamic culture under standard conditions, the experiment was performed with four different groups: 1) EGM2 served as control, 2) LPS was added to EGM-2 in a concentration of 100 ng/mL, 3) PBMCs were diluted in EGM2 with a final concentration of 1.5 × 10^6^ cells/mL, 4) For imitation of inflammatory conditions, LPS-activated PBMCs were added to the culture system at the same concentrations. After 24 h culture under experimental conditions, the culture was terminated by either fixation of the cells with ice-cold methanol (VWR) for immunocytochemistry (ICC) and scanning electron microscopy (SEM), cell detachment with accutase (Innovative Cell Technologies) for flow cytometry, or cell lysis and RNA isolation for quantitative polymerase chain reaction (qPCR) analysis.

### Leukocyte Adhesion Assay

For the leukocyte adhesion assay, PBMCs were stained with 5-chloromethylfluorescein diacetate (CMFDA, 1 µM, Thermo Fisher) prior to dynamic culture. After dynamic culture, microfluidic devices with HUVECs and attached PBMCs were fixed with ice-cold methanol for 10 min at -20 °C, and carefully washed three times with DPBS. Microscopy of the endothelial cells with attached PBMCs was performed (AxioObserver Z1, Carl Zeiss with AxioCam MRm with ZEN blue 3.6 software). Endothelialised area of each brightfield image was determined with GNU Image Manipulation Program software (GIMP V2.10.34, The GIMP team), and the number of attached PBMCs was calculated with open-source CellProfiler software V4.2.5 [20].

### Immunocytochemical Staining

The endothelialised membranes were carefully cut from the microfluidic devices, and ICC was performed with antibodies against CD31 and von Willebrand factor (vWf) and DAPI (Carl Roth) to counterstain nuclei as previously described [21]. For microscopy, a drop of mounting medium (Ibidi) was placed in a microscopy-suited µ-Slide chambered coverslip (Ibidi), and the sample was placed into the chamber with the cell-side down. Microscopy was performed with an inverted confocal laser scanning microscope (LSM 980 with Airyscan 2, Zeiss).

**Table 1:**
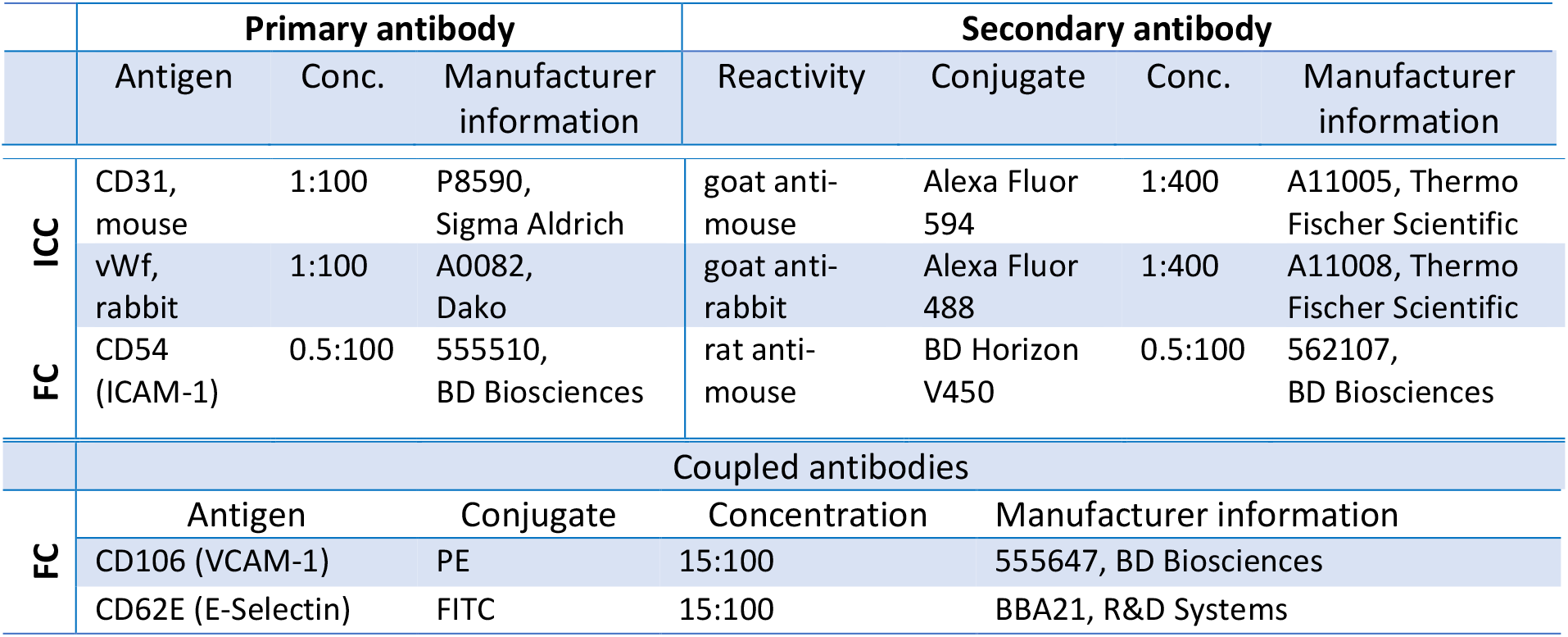
Antibodies used for immunocytochemistry (ICC) and flow cytometry (FC)

### Flow Cytometry Analysis

After dynamic culture, the cells were washed with DPBS, and treated with accutase for 3 minutes at 37 °C to ensure gentle cell detachment. After cell collection, live-dead staining (Zombie Aqua, BioLegend) of the cells was performed according to manufacturer’s instructions. After washing, flow cytometry antibody staining was performed according to the manufacturer’s instructions. The cells were resuspended in PBS with 1% FBS, and all measurements were performed with FACS Canto cytometer (BD Biosciences). Mean fluorescence intensities (MFI) of the adhesion molecule expression were evaluated and all data were processed using FlowJo software (V10.8.1, FlowJo LLC).

### Scanning Electron Microscopy

SEM analysis was performed by the facility for electron microscopy of the medical faculty of RWTH Aachen University. The methanol-fixed samples were transferred to 3% glutaraldehyde solution (Agar scientific) in 0.1 M sodium phosphate buffer (Merck). Dehydration was performed with ascending ethanol series (30%; 50%; 70%; 90%; 100%; 100%; 100%). Critical point drying in liquid CO_2_ (Polaron, Quorum Technologies Ltd) was performed before coating of the samples with a 10 nm gold/palladium layer (Sputter Coater EM SCD500, Leica). Scanning electron microscopy was performed in a high vacuum environment at 10 kV voltage (Quattro S, Thermo Fischer Scientific).

### Quantitative PCR Analysis

RNA was extracted from the cells after dynamic culture by removing the medium from the channels and adding RLT buffer (RNeasy Kit, Qiagen) for cell lysis and consequent RNA isolation. The cells were lyzed without separation of endothelial cells from potentially attached PBMCs in interest of time to process samples for RNA isolation. Collection of RNA from PBMCs was isolated from the cell-medium-suspension, such that no adherent endothelial cells were included. For RNA isolation from PBMCs, medium suspension from dynamic culture was centrifuged at 500 g for five minutes and the pellet containing PBMCs was lysed with the RLT buffer before RNA isolation according to the manufacturer’s protocol. Finally, RNA was eluted with RNAse-free water. RNA was reverse transcribed using a PrimeScript™ RT Reagent Kit (Takara Bio Europe), and PCR reactions were performed using iTaq Universal SYBR Green Supermix (Bio-Rad), according to the manufacturers’ protocols. The mRNA expression levels of VCAM-1, ICAM-1, E-selectin, IL-8, IL-6, IL-10, TNFα, MCP-1, TM, TPA, vWF, VE-cadherin, CD31, NOS3, EDN1 were measured using quantitative real-time PCR and normalized to the mRNA expression level of different reference genes. Supplementary Table S3 describes all used primers and annealing temperatures. CFX Maestro Software 1.1 (Bio-Rad) was used to determine the most stable reference genes: eukaryotic translation initiation factor 4A2 (EIF4A2) and ribosomal protein L13A (RPL13A). The term reference gene index (ref. gene index) was introduced as the normalization was always performed against these two genes. All PCR reactions were run on a CFX Connect Real-Time PCR Detection System (Bio-Rad) using the following protocol: 40 cycles of 10 s denaturation at 95 °C followed by 10 s annealing and 15 s amplification at 72 °C. The uncorrected RFU values using LinRegPCR version 2020.0 were used to determine the PCR efficiency [22]. The CFX Maestro Software 1.1 (Bio-Rad) was used for the relative quantification, and the evaluation algorithm was based on the ddCt method.

In order to distinguish between the effects coming from endothelial cells and PBMCs, principal component analysis (PCA) was performed based on the qPCR data in Python using the Scikit-learn module [23]. The standardization of the data was performed using the StandardScaler of Scikit-learn prior to PCA. Plotly Express (Plotly Technologies Inc.) was used for the graphical representation of the results. The samples were coloured depending on the culture conditions. The loadings of the genes were plotted as arrows with the length of the arrow representing the value of the loadings.

### Statistical Analysis

All experiments were performed with three independent biological replicates (*n* = 3). The results are presented in mean ± standard deviation. The data analysis was performed with Excel 2016 (Microsoft), and Prism 9 (9.4.0, Graphpad Software). Statistical evaluation of TGA and leukocyte adhesion assay was performed with Welch‘s *t*-test and Mann-Whitney test, respectively. Flow cytometry and qPCR data were analysed for the normal distribution using residual plots, Shapiro-Wilk and Kolmogorov-Smirnov tests as diagnostics. In case of normal distribution, ordinary one-way ANOVA with Tukey‘s multiple comparisons test was performed for the statistical analysis. A non-parametric Kruskal-Wallis test was performed if no normal distribution was confirmed (Shapiro-Wilk and Kolmogorov-Smirnov tests). A p-value below was considered statistically significant (labelled with *).

## Results

### Endothelial cells reduce thrombin generation on gas exchange membranes

To investigate if endothelial cells improve hemocompatibility of biofunctionalised PDMS gas exchange membranes, thrombin generation is measured (Figure 1). An increase in thrombin generation on RGD-coated PDMS is observed in contrast to endothelialised membranes, on which no thrombin generation occurs during the measurement time (Figure 1a). Thrombin generation is significantly reduced for endothelialised PDMS membranes compared to cell-free membranes, with the maximum thrombin generation of RGD-coated PDMS membrane reaching 2.8 × 10^3^ ± 8.8 × 10^2^ mU/(mL x min x cm^2^), whereas the confluent endothelial cell layer on the PDMS membrane does not induce relevant thrombin generation (Figure 1b).

**Figure 1.**
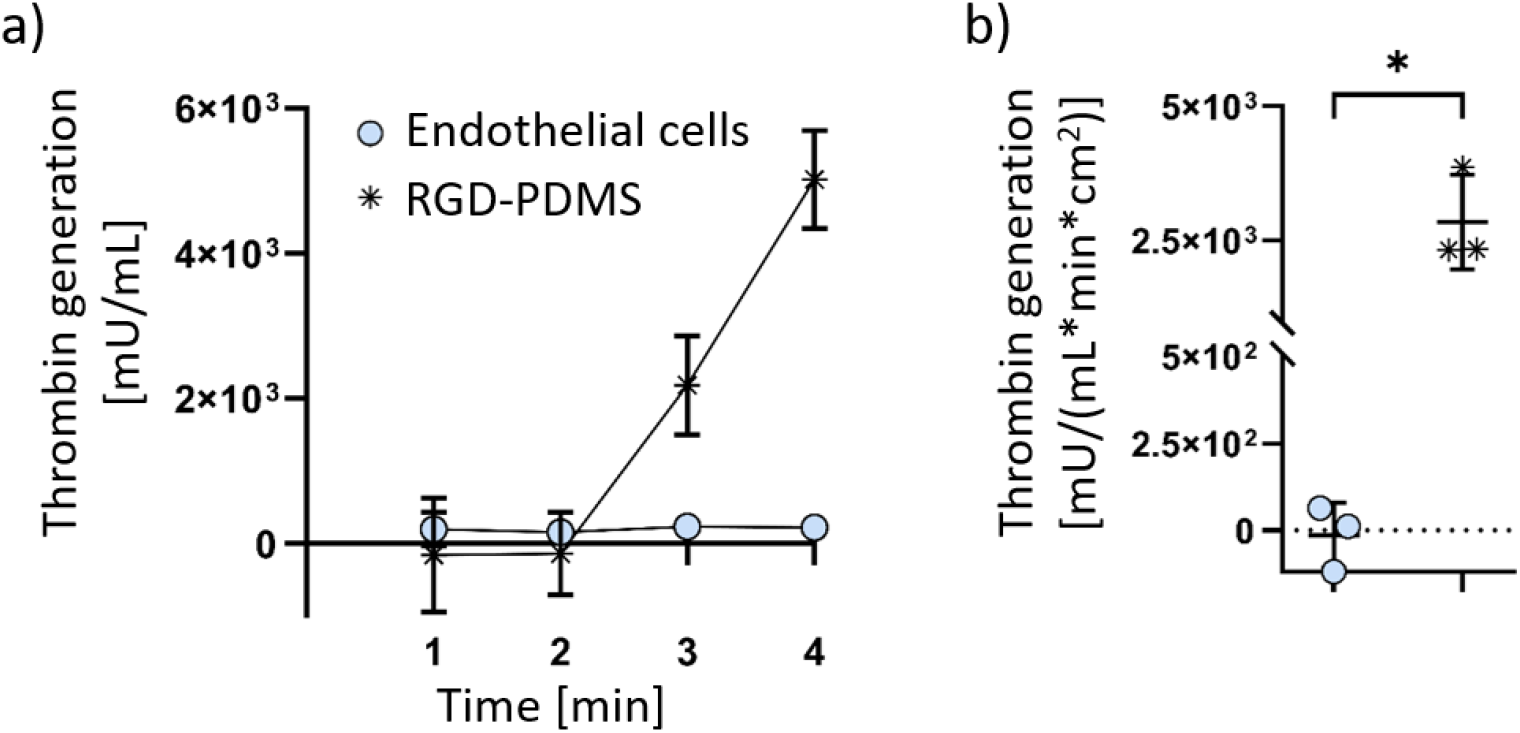
Thrombin generation on endothelialised PDMS membranes in comparison with non-endothelialised RGD-coated PDMS membranes. Thrombin generation over 4 minutes (a), and maximum thrombin generation during the assay (b) are shown. The data is shown as mean ± SD, with symbols in b) representing the individual data points. P-value below 0.05 was considered statistically significant (labelled with *).

### Inflammatory conditions disrupt the endothelial layer integrity

Maintenance of a confluent endothelial cell layer is essential for successful application of a biohybrid lung. ICC staining and SEM are performed to evaluate the ability of endothelial cells to withstand WSS and preserve cell layer confluence during dynamic culture under inflammatory conditions (Figure 2d, Figure 2h). Endothelial cells dynamically cultured with either only LPS or PBMCs or in standard cell culture medium are used as controls (Figure 2a-c, e-g).

**Figure 2.**
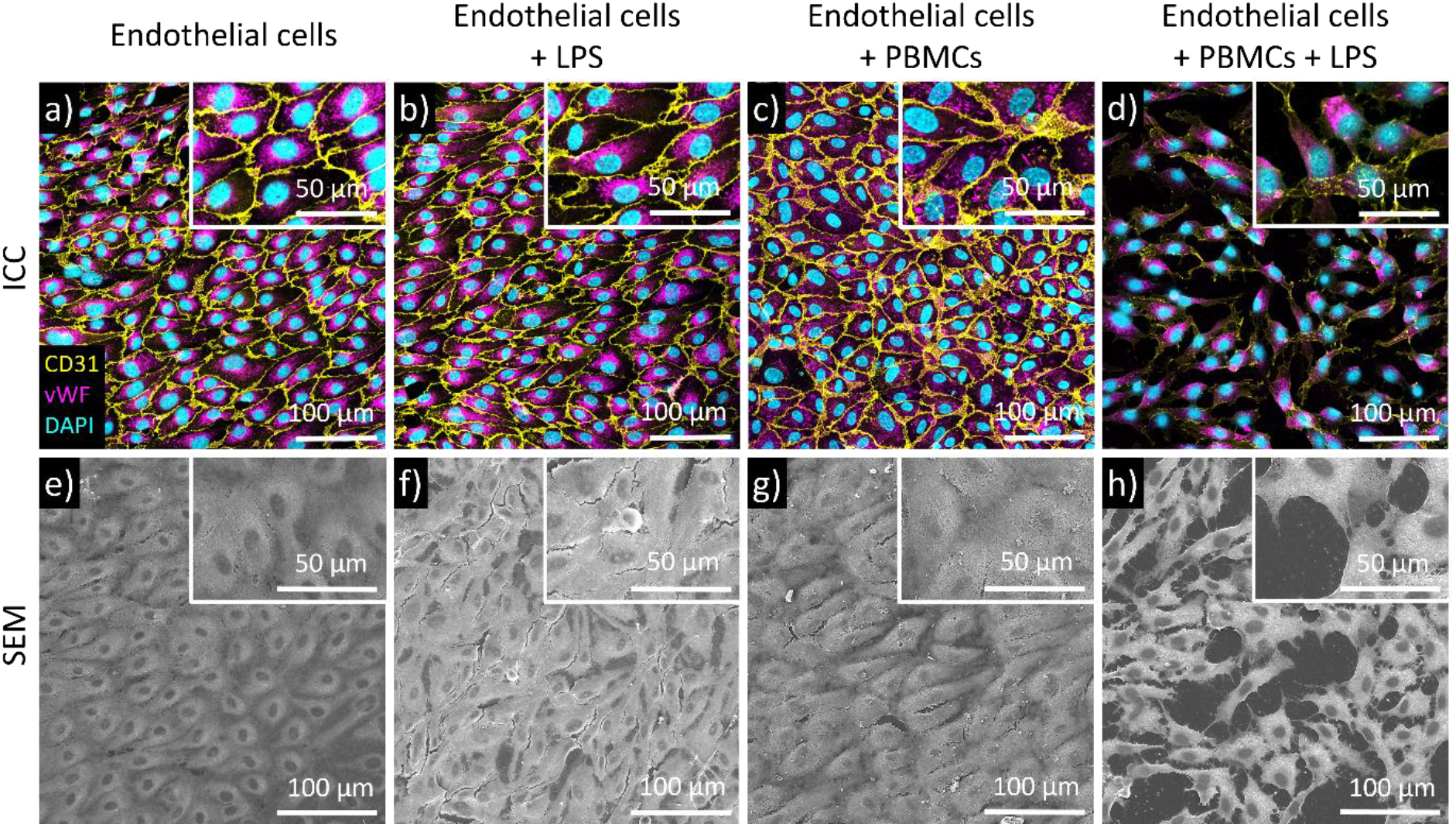
Evaluation of endothelial cell morphology. Immunocytochemical staining of endothelial cells against CD31 (yellow), vWF (purple), and DAPI (blue)-stained cell nuclei (a - d); scanning electron microscopy of endothelial cell layer after dynamic culture (e-h). Representative images of following conditions are shown: endothelial cells (medium control) (a, e); endothelial cells with LPS-treatment (b, f); endothelial cells cultured with PBMCs (c, g); endothelial cells cultured with LPS-treated PBMCs (d, h).

Cell-cell adhesions are shown by CD31 staining (yellow), while vWF is found in the cytoplasm (pink). Culture of endothelial cells with either LPS or PBMCs does not change their morphology or expression pattern of CD31 (Figure 2e-g, a-c, respectively). In contrast, a loss of cell confluence is observed upon culture of endothelial cells under inflammatory conditions. The endothelial cell layer appears disrupted (Figure 2d, h) and a substantial reduction of CD31-cell-cell adhesions is confirmed by antibody staining (Figure 2d). vWF expression does not seem to be altered by culture under inflammatory conditions (Figure 2d) in comparison to the controls (Figure 2a-c).

### Inflammatory conditions increase leukocyte adhesion

Leukocyte adhesion to endothelial cells is a key characteristic of inflammatory processes *in vivo*. In this work, it is directly and indirectly evaluated by measuring leukocyte adhesion (Figure 3a) and expression of adhesion molecules ICAM-1, E-Selectin, and VCAM-1 (Figure 3b-g) via flow cytometry, respectively.

**Figure 3.**
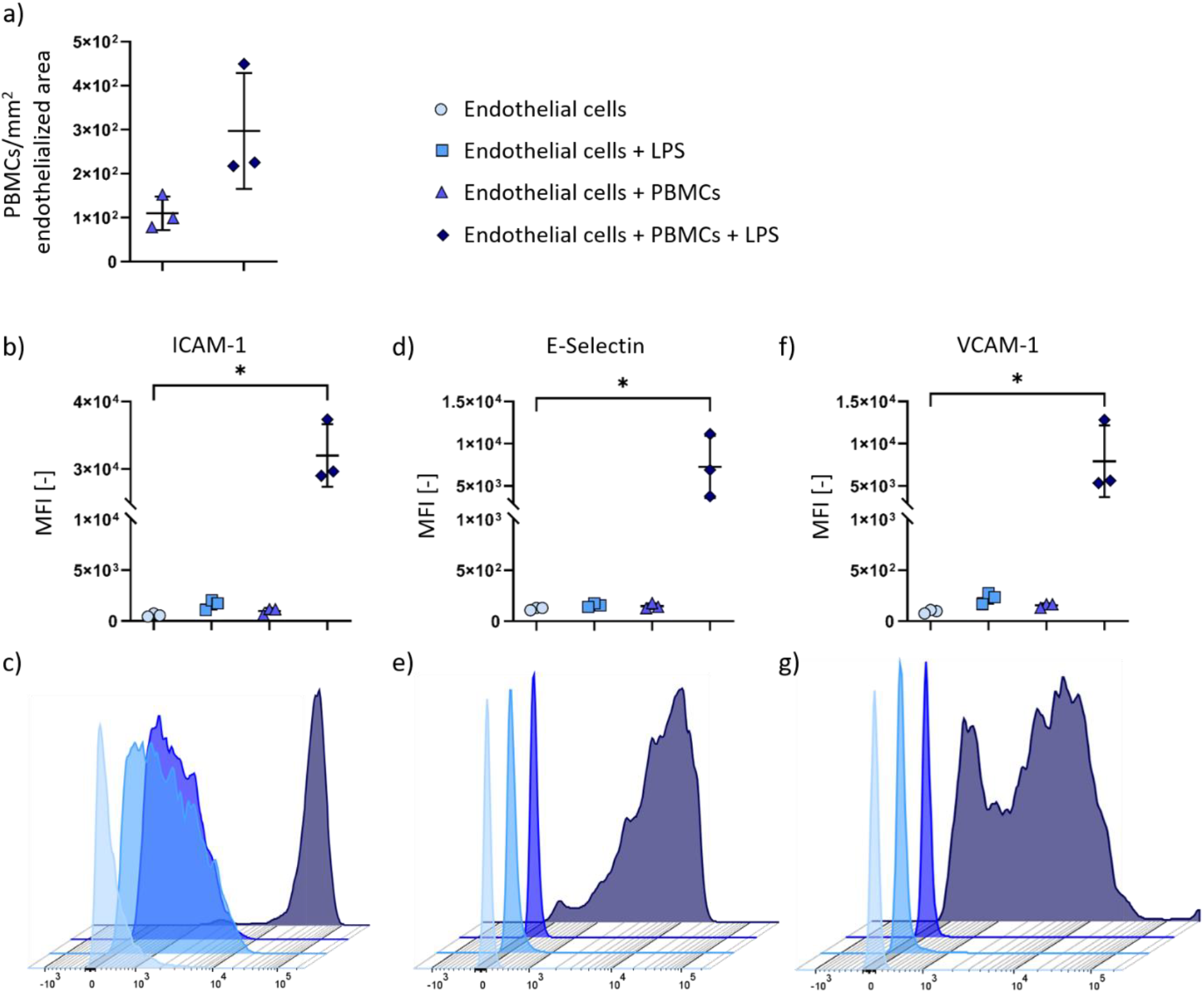
Leukocyte adhesion on endothelial cells (*n=3*) during dynamic culture under inflammatory conditions. Leukocyte adhesion assay (a) comparing PBMC-adhesion to endothelial cells with and without LPS-activation (*n=3*). Flow cytometry (b-g) of endothelial cells for following cell adhesion molecules: b), c): Intercellular adhesion molecule-1 (ICAM-1); d), e): E-Selectin; f), g): Vascular cell adhesion molecule 1 (VCAM-1). Mean fluorescent intensities (MFI) of adhesion molecules expression calculated from *n=3* independent donors are shown (b, d, f), as well as representative histograms of one donor (c, e, g). P-value below 0.05 was considered statistically significant (labelled with *).

Adhesion of PBMCs is increased on endothelial cells under LPS-treatment (Figure 3a). In detail, 110 ± 31 PBMCs/mm^2^ adhere to endothelial cells during LPS-free culture, and 297 ± 107 PBMCs/mm^2^ to endothelial cells with LPS-treatment. Representative pictures of endothelial cells treated with stained PBMCs, with and without LPS activation, are shown in Supplementary Figure S2. Flow cytometry reveals a significant increase in ICAM-1, E-Selectin, and VCAM-1 expression for culture of endothelial cells with LPS-activated PBMCs in comparison to the medium control (Figure 3b-g, see Supplementary table S2 for all MFI data).

### Inflammatory conditions induce the expression of pro-inflammatory proteins in endothelial cells

Endothelial cell behaviour during dynamic culture under inflammatory conditions is evaluated by quantitative mRNA analysis. The results are shown in Figure 4. Expression of the following genes is analysed to investigate inflammatory processes: ICAM-1, E-Selectin, VCAM-1, IL6, IL8 (gene CXCL8), IL10, TNFα, and MCP-1 (gene CCL2) (Figure 4a-h). The mRNA expression of endothelial adhesion molecules ICAM-1, VCAM-1, and E-Selectin is increased upon culture under inflammatory conditions, with a significant result for ICAM-1 expression compared to the control. Similarly, a higher expression of the pro-inflammatory genes IL6 and IL8, as well as the anti-inflammatory gene IL10 is observed in inflammatory conditions, along with a significant increase in TNFα and MPC-1 expression. No significant differences are observed in the expression of the following genes involved in thrombogenicity regulation in endothelial cells: vWF, TM (gene THBD), and TPA (gene PLAT) (Figure 4i, l, o). NOS3 gene expression is significantly reduced, while its antagonist, EDN1, shows a significant increase in endothelial cells cultured under inflammatory conditions (Figure 4j-k). No change in VE-Cadherin (gene CDH5) expression is observed (Figure 4m). Expression of CD31 (gene PECAM-1), is significantly decreased in inflammatory conditions (Figure 4n).

**Figure 4.**
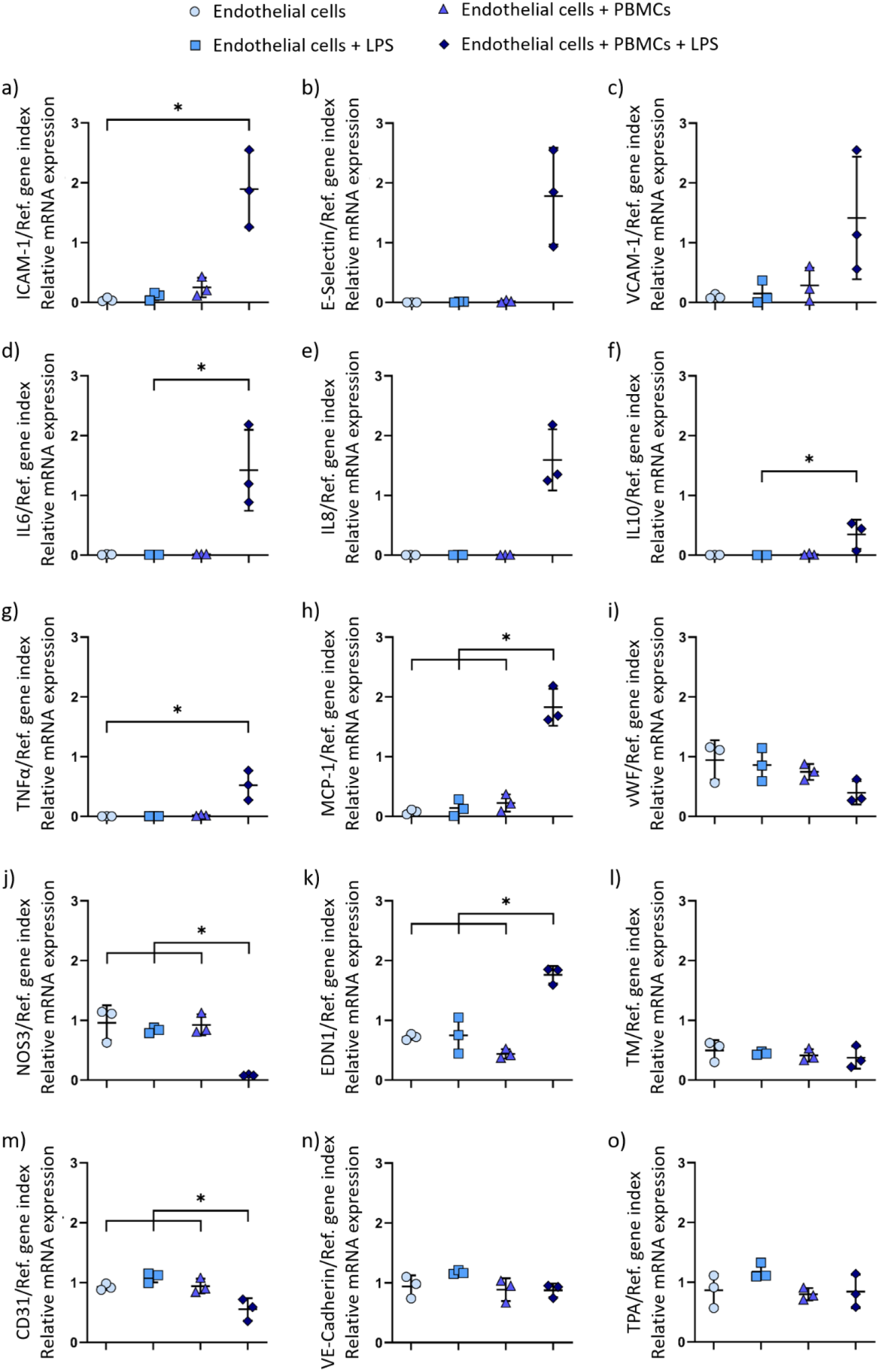
Relative mRNA expression in endothelial cells (*n=3*) after dynamic culture under inflammatory conditions (endothelial cells + PBMCs + LPS), with following groups as controls: endothelial cells, endothelial cells + LPS, endothelial cells + PBMCs. Endothelial cells were analysed for relative mRNA expression of a) Intercellular adhesion molecule-1 (ICAM-1); b) E-Selectin; c) Vascular cell adhesion molecule 1 (VCAM-1); d) Interleukin 6 (IL6); e) Interleukin 8 (IL8); f) Interleukin 10 (IL10); g) Tumor necrosis factor alpha (TNFα); h) Monocyte chemoattractant protein-1 (MCP-1); i) Von Willebrand factor (vWF); j) Nitric oxide synthase 3 (NOS3); k) Endothelin 1 (EDN1); l) Thrombomodulin (TM); m) Platelet and endothelial cell adhesion molecule 1 (PECAM-1, CD31); n) Vascular endothelial cadherin (VE-Cadherin); o) Tissue plasminogen activator (TPA) in relation to a reference gene index consisting of EIF4A2 and RPL13A. Data are shown as mean ± SD with symbols representing the individual data points. P-value below 0.05 was considered statistically significant (labelled with *).

A multidimensional PCA was performed to further evaluate qPCR data (Figure 5). PCA showed a clear separation of endothelial cells dynamically cultured under inflammatory conditions from all control groups (Figure 5a). Furthermore, clear separation of PBMCs from all endothelial cell groups was observed, with an additional separation between the PBMCs and LPS-activated PBMCs (Figure 5a). Three groups of genes can be identified considering the loadings of the genes (Figure 5b). While the mRNA expression of ICAM-1, VCAM-1, E-Selectin, IL6, IL8, MCP-1, and EDN1 affect the grouping of endothelial cells cultured under inflammatory conditions, mRNA expression of VE-Cadherin, CD31, vWF, and NOS3 genes appears to be relevant for endothelial cells cultured under all other conditions. PBMCs (with and without LPS) strongly express TNFα and IL10 mRNA (Figure 5b).

**Figure 5.**
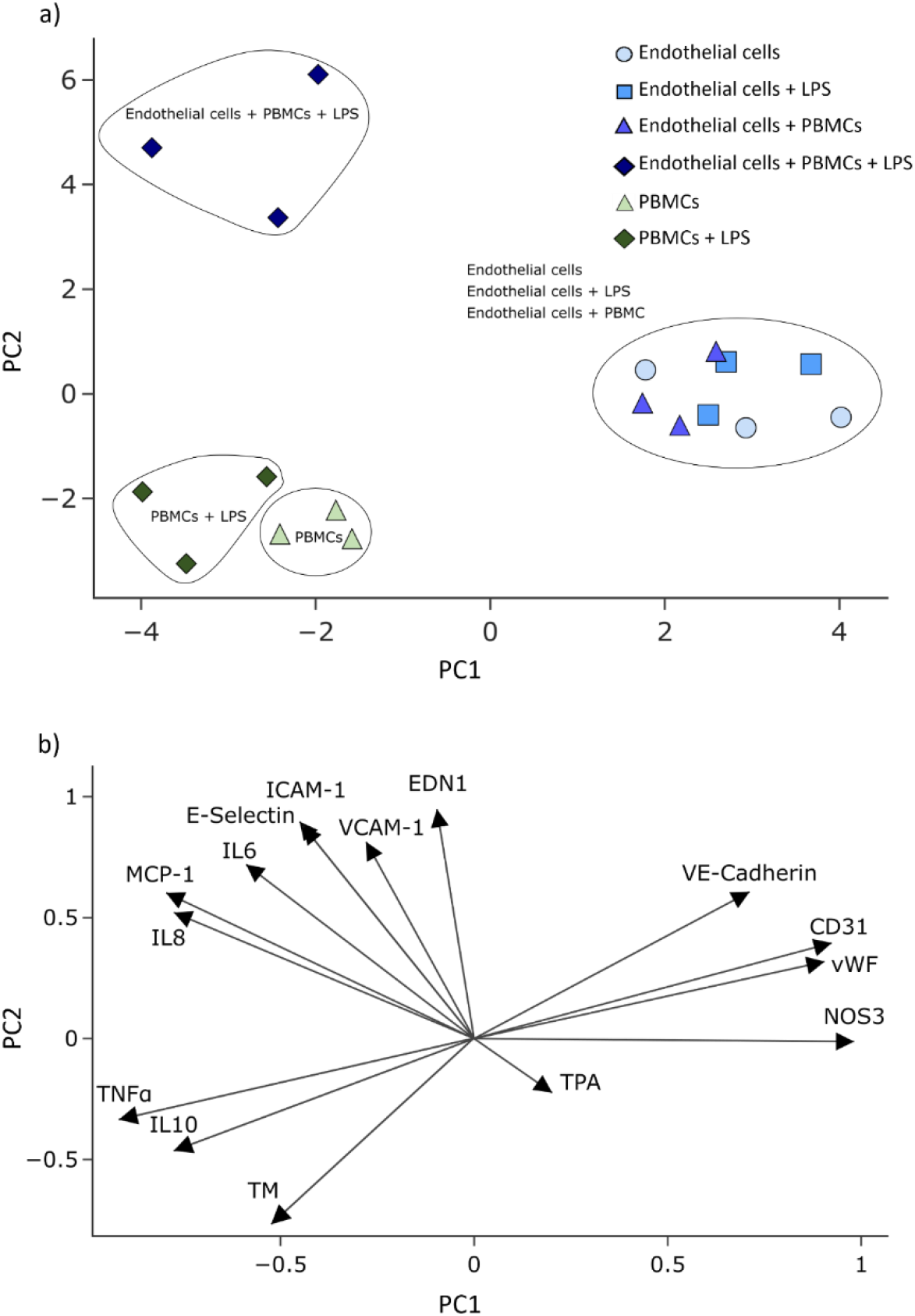
Principal component analysis of mRNA expression data. Results are presented as scatter plots where each symbol represents one sample. Samples are coloured for the different culture conditions (a). The loadings of investigated genes are plotted as arrows (b).

## Discussion

In the current study, we highlight the importance of replicating physiological conditions closely *in vitro* to advance the translational potential of biohybrid medical devices. Specifically, endothelialisation of gas exchange membranes has been proposed in several studies as a possible solution to overcome hemocompatibility issues of ECMO devices, mainly by proof-of-concept studies as well as methodologies for endothelialisation and the evaluation of the ability of endothelial cells to maintain confluence and functionality during long-term dynamic culture under non-inflammatory conditions [4, 6, 24].

We measured thrombin generation to evaluate if the thrombogenicity of PDMS membranes improves by endothelialisation. The result supports the general hypothesis that endothelialisation of artificial membranes reduces membrane thrombogenicity and thus improves the devices’ hemocompatibility. We assume that the thrombogenicity of endothelialised membrane remains significantly lower than that of uncoated membranes, as long as the endothelial cell layer is fully confluent and does not expose the membrane surface underneath it [25].

An inflammatory status has been reported in many patients whose condition could be treated with a biohybrid device such as endothelialzed ECMO, or a vascular graft [10, 26, 27]. Systemic inflammatory response is characterized by highly elevated pro- and anti-inflammatory cytokine levels [28]. Endothelial cells respond to inflammation with endothelial activation which eventually results in increased leukocyte adhesion, decreased barrier function, vasoconstriction, and vascular leakage and changes in thromboresistance [13]. This inflammation-induced vascular dysfunction is reversible with appropriate treatment *in vivo* [29]. To the authors’ knowledge, no investigation of endothelial cells on artificial membranes has been performed under inflammatory conditions. Thus, we used a dynamic microfluidic culture system that allows for dynamic culture of endothelial cells on gas exchange membranes under inflammatory conditions imitated by the addition of LPS-activated PBMCs. Essential differences have been previously observed in the endothelial cell response to cytokine induction in static versus dynamic culture conditions, indicating the importance of evaluating the endothelial cell response to inflammation under dynamic conditions [30].

Evaluation of endothelial cell layer after inflammatory treatment showed a loss of endothelial cell layer confluence. CD31 expression significantly decreased in endothelial cells under inflammatory conditions on mRNA and protein level. CD31 regulates vascular permeability, as well as inhibits the activation of circulating thrombocytes and leukocytes [31]. A combination of cytokines TNFα and interferon-γ (IFNγ) has been reported to decrease CD31 expression [32]. Consistent with these findings, we observed upregulation of TNFα gene expression in endothelial cells after inflammatory treatment. Notably, endothelial cells and PBMCs can express TNFα [33, 34]. According to the PCA of qPCR results, the TNFα gene expression primarily occurs in LPS-activated PBMCs. Thus, we assume that the reduction in CD31 expression in endothelial cells occurs due to increased TNFα expression in LPS-activated PBMCs.

*In vivo*, leukocyte adhesion is an essential part of the defence mechanisms during inflammation. After activation, leukocytes tend to accumulate around the site of inflammation [35]. Leukocytes start a rolling movement along the endothelial cells, attaching to the endothelial cell adhesion molecule E-Selectin [36]. ICAM-1 and VCAM-1, as ligands for lymphocyte function adhesion molecule-1 (LFA-1) and very late activation antigen-4 (VLA-4), respectively, allow leukocytes to firmly adhere to the endothelial cells [37]. Following, leukocytes transmigrate through the endothelial cell layer towards the inflamed tissue. Although a necessary and reversible process during healthy vascular physiology, leukocyte transmigration can result in vascular leak, expanding permeability of endothelial cell layer in pathological conditions [38]. Consistent with the *in vivo* reaction of endothelial cells to inflammation, we observed an increase in endothelial cell adhesion molecules E-Selectin, ICAM-1 and VCAM-1 on protein and gene expression level. These results confirm endothelial cell activation resulting in increased leukocyte adhesion during culture under inflammatory conditions. We assume that the lack of underlying tissue further promotes endothelial cell detachment and restricts the endothelial cells from restoring the confluence.

Because cyto- and chemokines are an important mechanism for intercellular communication, we evaluated the gene expression of several pro- and anti-inflammatory factors. Upregulation of IL6, IL8, and MCP-1 gene expression in endothelial cells under inflammatory conditions confirms cell activation. This is associated with vascular leakage under septic conditions, which is consistent with our results [39]. Additionally, MCP-1 is responsible for attracting leukocytes to the site of injury [40]. In contrast, IL10 acts as an anti-inflammatory cytokine. IL10 has been shown to inhibit pro-inflammatory cytokine production, i.e. TNFα and IFNγ, and support wound healing and tissue repair [41, 42]. In our study, a significant increase in IL10 gene expression was detected under inflammatory conditions. PCA revealed that IL10, similar to TNFα, is mostly expressed by PBMCs and not endothelial cells in the microfluidic system. Thus, we assume that the observed upregulation of IL10 comes as an effect from the RNA being isolated from endothelial cells with the attached PBMCs. Altogether, IL10 expressed by PBMCs might be a potential treatment target to decrease pro-inflammatory cytokine expression and promote endothelial cell layer recovery.

NOS3 is responsible for nitric oxide (NO) production, and thereby reduces thrombocyte reactivity and thrombogenicity of endothelial cells [43]. The reduction in NOS3 gene expression in endothelial cells under inflammatory conditions could lead to a decrease in NO production. In a biohybrid lung setup, this in turn could lead to an increased thrombosis risk in patients with inflammation. NOS3 protects against systemic inflammation by reducing ICAM-1 and IL6 expression, which makes it an interesting potential therapeutic target for inflammation treatment [44]. During inflammation, the disruption of vascular homeostasis leads to upregulation of EDN1, which acts as an antagonist to NOS3 [45]. This has been confirmed by our investigation of EDN1 gene expression. LPS-induced sepsis is associated with increased EDN1 expression [46]. In endothelial cells, EDN1 binds to endothelin receptor type B (ETB) that leads to an increased NOS3 activation, and in turn caused NO-induced vasodilation [47]. In general, reduced expression of NOS3 and increased expression of EDN1 in endothelial cells during inflammation points to endothelial cell dysfunction due to KLF2 expression destabilisation [48].

The study represents the first investigation of endothelialised membranes for potential application in a biohybrid device model under inflammatory conditions. We deem this essential before proceeding with (pre-) clinical studies to prove the feasibility of biohybrid devices. Our study emphasizes the importance of (biohybrid) device testing in conditions similar to later application in patients. A condition such as inflammation can drastically influence the potential device performance, as shown in this study.

In the setup used for this study, the reactions of endothelial cells to complex inflammatory stimuli provided by the combination of PBMCs and LPS very closely mimic the reactions observed *in vivo*. The setup is highly flexible: the culture substrate can easily be exchanged, wall shear stresses can be varied, and immune blood cell or LPS concentration can be modified as well as the used cell types changed, e.g. by adding granulocytes or thrombocytes. Its use extends to a versatile disease model not only for the study of endothelial behaviour in sepsis but for the complex interactions between endothelial cells and the immune system.

## Conclusion

The presented setup allowed us to investigate endothelial cell behaviour on artificial membranes under inflammatory conditions. Exposure to LPS-activated PBMCs causes loss of endothelial cell layer confluence, as well as increased leukocyte adhesion, accompanied by upregulation of inflammation-associated genes in endothelial cells. The observed loss of endothelial cell layer integrity puts a big question mark on the concept of biohybrid device endothelialisation under inflammatory conditions. An endothelial layer failing already after 24 h would expose the underlying artificial surface, triggering thrombus formation and negating the intended benefits. This work highlights the need for careful consideration of patient inflammatory status in the design and evaluation of biohybrid devices. Future studies should incorporate such translational insights to refine device strategies, ensuring their efficacy and safety in clinical application.

## List of Abbreviations

ABM: antibiotic-antimycotic solution
CD31: Cluster of differentiation 31
CMFDA: 5-Chloromethylfluorescein diacetate
COPD: Chronic obstructive pulmonary disease
DAPI: 4’,6-Diamidino-2-phenylindole
DPBS: Dulbecco’s phosphate buffered saline
ECMO: Extracorporeal membrane oxygenation
EIF4A2: Eukaryotic translation initiation factor 4A2
EDN1: Endothelin 1
EGM2: Endothelial growth medium 2
ETB: Endothelin receptor type B
FACS: Fluorescence-activated cell sorting
ICAM-1: Intercellular adhesion molecule 1
IFNγ: Interferon gamma
IL-6: Interleukin 6
IL-8: Interleukin 8
IL-10: Interleukin 10
KLF2: Krüppel-like factor 2
LFA-1: Lymphocyte function-associated antigen 1
LPS: Lipopolysaccharide
MCP-1: Monocyte chemoattractant protein 1
MFI: Mean fluorescence intensity
NO: Nitric oxide
NOS3: Nitric oxide synthase 3
PBMCs: Peripheral blood mononuclear cells
PCA: Principal component analysis
PECAM-1: Platelet endothelial cell adhesion molecule 1
PDMS: Polydimethylsiloxane
qPCR: Quantitative polymerase chain reaction
RGD: Arginine-Glycine-Aspartate
RNA: Ribonucleic acid
RFU: Relative fluorescence units
RPPL13A: Ribosomal protein L13A
SEM: Scanning electron microscopy
SIRS: Systemic inflammatory response syndrome
TGA: Thrombin generation assay
TNFα: Tumor necrosis factor alpha
TM: Thrombomodulin
TPA: Tissue plasminogen activator
VE-Cadherin: Vascular endothelial cadherin
vWf: von Willebrand factor
VCAM-1: Vascular cell adhesion molecule 1
VLA-4: Very late antigen 4
WSS: Wall shear stress;

## Acknowledgements

This research was funded by the German Research Foundation (Deutsche Forschungsgemeinschaft, DFG) in the framework of the priority program “Towards an Implantable Lung” (grant numbers 313779459, 447717028, 447712946 and Start-Funding for Young Female Researchers) and within the Research Training Group 2415 (Project number: 363055819). The Scanning Electron Microscope Quattro S was funded by the DFG under grant number 495328185.

The authors would like to thank the blood donors and the Clinic for Gynaecology and Obstetrics, RWTH Aachen University Hospital, headed by Univ.-Prof. Dr. med. Elmar Stickeler for providing umbilical cords for human umbilical cord endothelial cells isolation. This work was supported by the Confocal Microscopy Facility as well as by the Flow Cytometry Facility, both core facilities of the Interdisciplinary Center for Clinical Research (IZKF) Aachen within the Faculty of Medicine at RWTH Aachen University. We thank Nathalie Steinke for excellent technical support. Supplementary Figure 1 and the graphical abstract were created with Biorender.com.

## Ethics Approval Statement

The study was conducted in accordance with the Declaration of Helsinki and approved by the Ethics Committee of the Medical Faculty of RWTH Aachen University (protocol code EK 22-390 and cBMB project number 323). All participants (or their authorized representative) provided informed consent.

## Data availability

The data that support the findings of this study are available from the corresponding author upon reasonable request.

## Author contributions

Conceptualization: A.L.T., C.G.C., K.T. and S.J; Formal analysis: M.C. and A.B.; Funding acquisition: K.T., C.G.C., A.L.T. and S.J.; Investigation: M.C., S.K., A.P., S.R. and A.B.; Methodology: M.C., S.K., A.P., S.R. and A.B.; Project administration: C.G.C., A.L.T., K.T. and S.J.; Resources: S.J.; Supervision: K.O., K.T., A.L., C.G.C., A.L.T., S.J.; Validation: M.C., A.B., A.L.T. and S.J., Visualization: M.C. and A.B., Writing – original draft: M.C.; Writing – review & editing: M.C., A.B., C.T.N., S.K., A.P., K.O., K.T., S.R., A.L., C.G.C., A.L.T. and S.J.

## Conflict of Interest Statement

The authors declare no competing interests or conflict of interest. The funders had no role in the design of the study; in the collection, analyses or interpretation of data; in the writing of the manuscript; or in the decision to publish the results.

## Notes

### Competing Interest Statement

The authors have declared no competing interest.

### Summary of Updates

Updated manuscript text: Uptadet abstract to shiift the focus of the manuscript towards more "Biohybrid device" topic

