## Supplemental files for "Modelling Inflammatory Endothelial Dysfunction: A Human In Vitro Platform for Translational Research"

**Supplementary material**

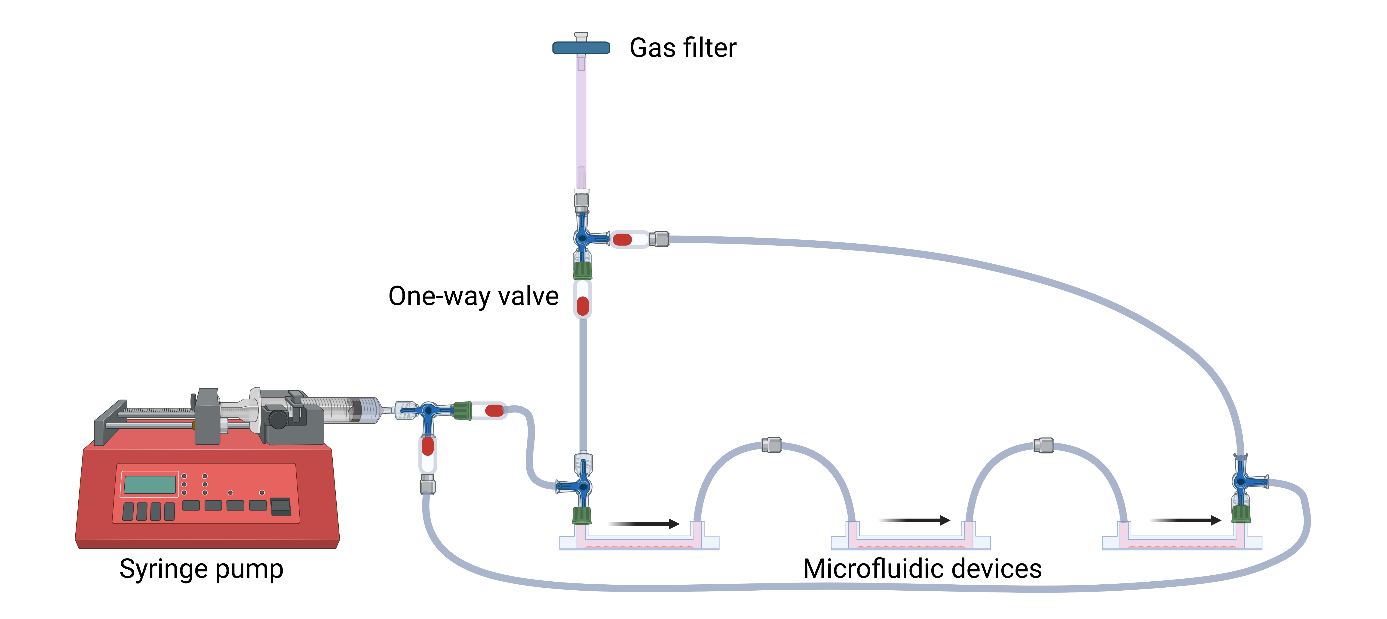

**Figure S1.** Bioreactor system with syringe pump and connected microfluidic devices. The bioreactor system was assembled using a syringe pump (LA-120, Landgraf Laborsysteme GmbH), syringe (B.Braun), gas filter (Merck), three-way valves (B.Braun), one-way valves (CODAN Medizinische Geräte GmbH) and Heidelberger extensions (B.Braun). Including one-way valves allowed for one-directional flow inside the microfluidic chambers during the syringe's pushing and pulling phase. The bioreactor system was assembled and cultured with EGM2 medium with 1% ABM at 37 °C and 5% CO_2_ 24 h prior to connection of microfluidic devices for system equilibration to reduce the amount of air bubbles in the tubing. Created in BioRender. Cheremkhina, M. (2025) https://BioRender.com/9q2cngd

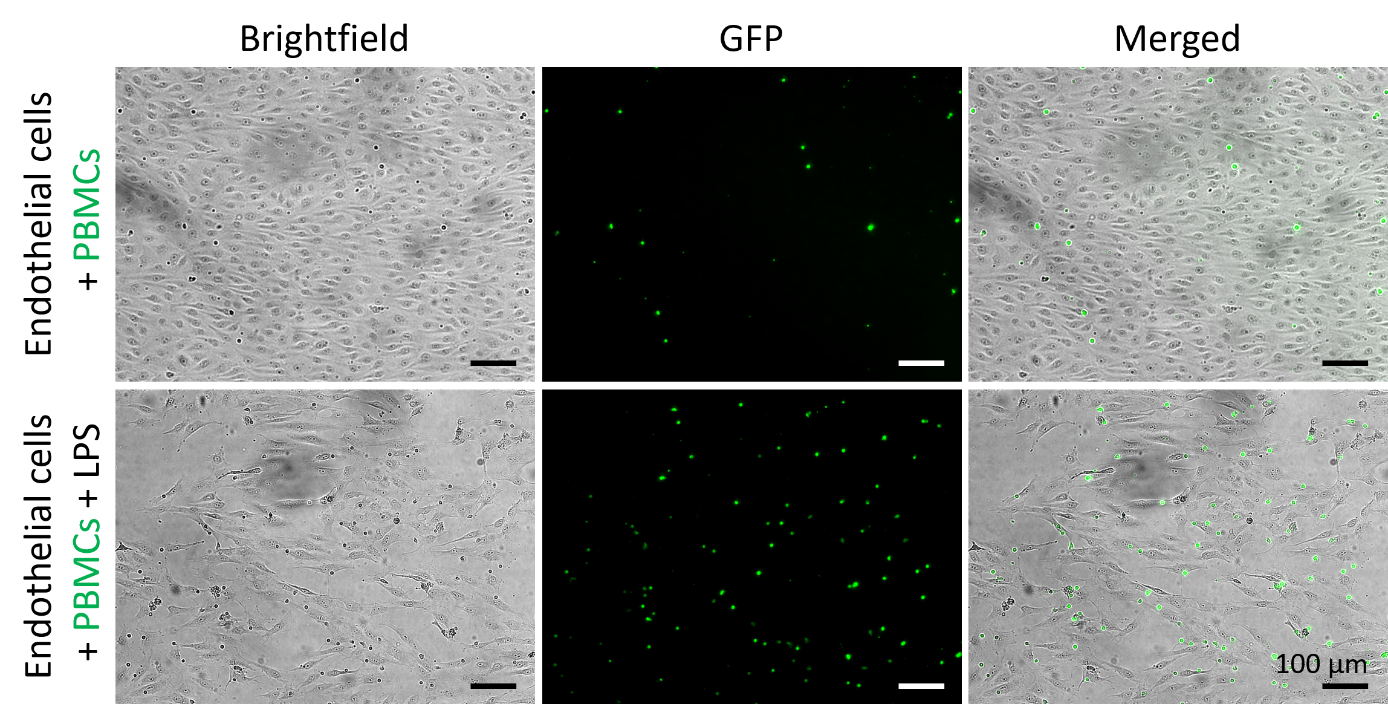

**Figure S2.** Representative images of leukocyte adhesion assay. PBMCs (green) dynamically cultivated on endothelial cells (brightfield) with or without LPS-activation. Scale bar: 100 µm.

**Table S1.** Percentage of positive HUVECs (n=4) stained for four endothelial cell-characteristic markers (CD31, CD105, CD144, and CD146) and one mesenchymal stromal cell-characteristic marker (CD90).

|  | *Percentage of positive cells  Mean ± SD [%]* |
| --- | --- |
| CD31 (PECAM-1) | 100 ± 0 |
| CD105 (Endoglin) | 99.98 ± 0.05 |
| CD144 (VE-cadherin) | 98.03 ± 1.26 |
| CD146 (MCAM) | 91.53 ± 15.63 |
| CD90 | 3.08 ± 1.84 |

**Table S2**. Mean Fluorescence Intensity (MFI) of ICAM-1, E-Selectin, and VCAM-1 expression measured by flow cytometry of endothelial cells (n=3) under the following conditions: no treatment, treatment with LPS, treatment with PBMCs, and treatment with LPS-activated PBMCs.

| *Adhesion molecule* | *Condition* | *MFI  Mean ± SD [-]* |
| --- | --- | --- |
| ICAM-1 | Endothelial cells | 569.7 ± 122 |
|  | Endothelial cells + LPS | 1607.3 ± 396 |
|  | Endothelial cells + PBMCs | 972 ± 249.7 |
|  | Endothelial cells + PBMCs + LPS | 31994.7 ± 3795.8 |
| E-Selectin | Endothelial cells | 122 ± 9.9 |
|  | Endothelial cells + LPS | 155.3 ± 14.7 |
|  | Endothelial cells + PBMCs | 148 ± 21.2 |
|  | Endothelial cells + PBMCs + LPS | 7256.3 ± 3024.3 |
| VCAM-1 | Endothelial cells | 94 ± 14.3 |
|  | Endothelial cells + LPS | 226.3 ± 44.2 |
|  | Endothelial cells + PBMCs | 154 ± 16.3 |
|  | Endothelial cells + PBMCs + LPS | 7918.7 ± 3465.4 |

**Table S3.** Primers used for qPCR and their annealing temperatures.

| *Gene* | *Sequence* | *Annealing temperature* |
| --- | --- | --- |
| ICAM-1 | forward: GGAGCCCGCTGAGGTCACGA | 66 °C |
|  | reverse: CGCTGGCAGGACAAAGGTCTGG |  |
| E-Selectin | forward: TTGCCCTATGCTACACAG | 56 °C |
|  | reverse: TTGAGTCCACTGAAGCCA |  |
| VCAM-1 | forward: GCAAGTCTACATATCACCC | 57 °C |
|  | reverse: AATCTTCCATCCTCATAGCA |  |
| IL6 | forward: GTGTGAAAGCAGCAAAGAG | 57 °C |
|  | reverse: AAGTCTCCTCATTGAATCCA |  |
| IL8 (gene CXCL8) | forward: GACATACTCCAAACCTTTCC | 60 °C |
|  | reverse: AACTTCTCCACAACCCTC |  |
| IL10 | forward: GCTGTCATCGATTTCTTC | 60 °C |
|  | reverse: GTCAAACTCACTCATGGC |  |
| TNFα | forward: TGAGCACTGAAAGCATGATCC | 60 °C |
|  | reverse: CGAGAAGATGATCTGACTGCC |  |
| MCP-1 (gene CCL2) | forward: ATGAAAGTCTCTGCCGCC | 57 °C |
|  | reverse: CTTCTTTGGGACACTTGCT |  |
| vWF | forward: TACCACAACCACCTGCCT | 57 °C |
|  | reverse: GTAAGTGAAGCCCGACCGA |  |
| TM (gene THBD) | forward: AGAGAAGAGACAAACACCT | 57 °C |
|  | reverse: TCCACAAGACCAGTAGAG |  |
| TPA (gene PLAT) | forward: TGCTACTTTGGGAATGGG | 57 °C |
|  | reverse: GTTCTGTGCTGTGTAAACCT |  |
| NOS3 | forward: CGAGTGAACGCGACAATCCT | 60 °C |
|  | reverse: GCTGCAAAGCTCTCTCCATTC |  |
| EDN1 | forward: CCTAAGACAAACCAGGTCGG | 60 °C |
|  | reverse: CTTTGCCAGTCAGGAACCA |  |
| VE-Cadherin (gene CDH5) | forward: TCAAGCGTGAGTCCGCAAGAA | 60 °C |
|  | reverse: AATGACAGCAGTGAGGTGGT |  |
| CD31 (gene PECAM-1) | forward: CAGCCAACTTCACCATCC | 57 °C |
|  | reverse: GAGAGCATTTCACATACGAC |  |
| EIF4A2 | forward: CCAAAAGGTAATTCTGGCACTTG | 60 °C |
|  | reverse: CGGGTGTACCAACAACAATATGT |  |
| RPL13A | forward: GCCCTACGACAAGAAAAGCG | 60 °C |
|  | reverse: TACTTCCAGCCAACCTCGTGA |  |
